# Analysing and identifying miRNAs from RNA-seq data using miRDeep2 tool in Galaxy, a practical guide

**DOI:** 10.1101/2021.10.19.464446

**Authors:** M. Rifqi Rafsanjani

## Abstract

Micro-RNAs (miRNAs) analysis from RNA-seq experiment data provides additional depth into the cellular gene regulation. Such analysis has been simplified using miRDeep2 tool available in freely accessible Galaxy Europe server and miRBase database that compiled the miRNAs in many organisms. Here, we are describing a step by step protocol on how to ultilised the tool and most importantly on how to prepared and compiled miRDeep2 results to be used for downstream analysis such as investigating the differential expression of the detected miRNAs. Currently miRDeep2 miRNAs count result output in the Galaxy are in the broken HTML page and processing such data are troublesome for further analysis unless the user setting up their software dependencies for the tool to run. Hence, we proposed a method to process this output so that it is usable for downstream processing without a single coding required.

## 1 Introduction

Micro-RNAs (miRNAs) are the non-coding RNAs that have many roles in regulating gene expression. Analysis of this miRNAs especially from the RNA-seq data could provide insights on how gene expression is regulated and with many powerful bioinformatic tools, miRNAs analysis could never been easier. In the free to use Galaxy Europe server^1^, several tools allows the analysis and identification of miRNAs such as PIPmiR PIPELINE (for plant miRNAs), miRanda (to find the potential targets site of miRNAs in the genomic sequence) and miRDeep2 (quantification of reads mapping to known miRBase precursors). With the abundance of RNA-seq data freely available from the online database, such identification of miRNAs opens up new possibilities for other researchers to use for their own analysis and application.

miRDeep2^2,3^, which was originally published in 2011 has identified many miRNAs in a wide variety number of organisms^2,4^, identified miRNAs in the exosomes^5^, used in toxicogenomics study^6^ and many other applications. However, the detailed step by step methodology on how to prepare the input and analyse the output has raised many questions in many research forums even to this date. Although miRDeep2 analysis is very straightforward, a clear guidance of how to obtain the inputs and manipulate the outputs are among of the minor obstacle especially for the early stage researchers. To resolve this particular issue, we attempt to describe a clear guide step by step protocol on how to do so with no coding experience required.

Current results from miRDeep2 Quantifier analysis in Galaxy Europe server contains a broken formatting of HTML file that summaries the miRNAs findings. This is good enough if the user would like to represent the findings in the available table form but for the further downstream analysis, a proper data compiling and structured is certainly needed. This is one of the issue we encountered using the tool hence the main objective of why this protocol is written.

### 1.1 Overview of the method

Prior to the analysis, user is expected to already registered to the Galaxy Server account, and already has RNA-seq cleaned reads prepared in the history panel. For on how to prepare the RNA-seq data, please refer to the Galaxy Training webpage^7^. Other inputs used in this guide will be mentioned on how to obtained and processed it along the way. Additional software required for this particular guide are summaries in Section 2.1. Overview of this guide are summaries in the Figure 1.

**Figure 1:**
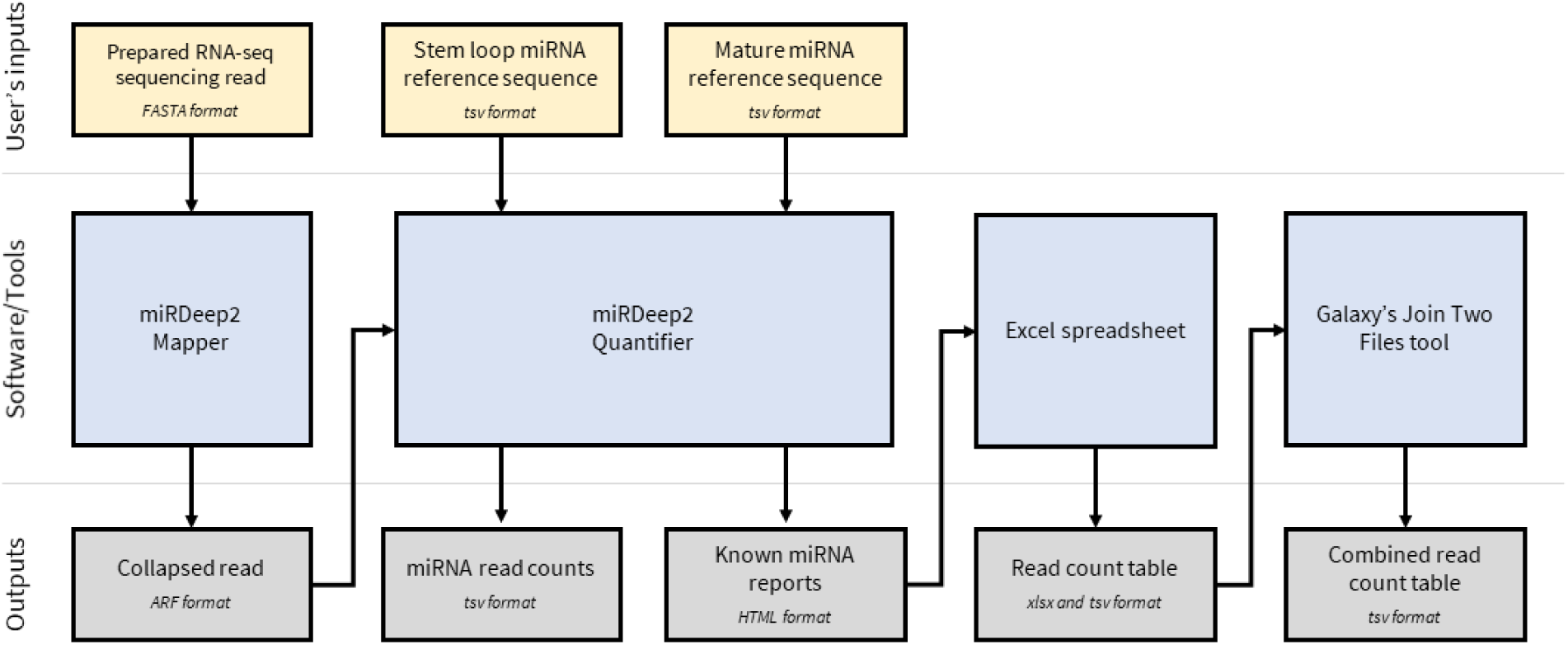
Overview of the guide. The flowchart diagram shows the overview of the step of this guide to analyse and process the miRDeep2 inputs and outputs.

### 1.2 Limitations

This guide only attempt to only describe the processing part of miRDeep2 and the formatting of the ouputs, so the input sequence of cleaned RNA-seq data is assumed to be prepared by the user. The final data obtained from this guide can be use for downstream processing such as in edgeR or limma tool in Galaxy and processing the input for these tools are also described on how to compiled the processed miRNAs read counts.

## 2 Procedure

### 2.1 Materials and Equipment

- A computer with an internet access
- Galaxy Europe registered account
- Prepared and trimmed RNA-seq data in FASTA format
- Microsoft Office Excel or any other spreadsheet software
- Notepad or any other text editor software
- Access to tools used in the Galaxy: mirDeep2 Mapper, mirDeep2 Quantifier and Join Two Files.

### 2.2 Methods

#### 2.2.1 Mapping the RNA-seq reads using miRDeep2 Mapper

miRDeep2 can be accessed from the tools panel at the Galaxy Europe website (https://usegalaxy.eu/) by searching the tool name into the search box. You will get three results which are miRDeep2 package, miRDeep2 Quantifier and miRDeep2 Mapper. We will map our RNA-seq reads using miRDeep2 Mapper first by choosing your uploaded sequencing reads in FASTA format.

1. Open miRDeep2 Mapper and key in the following parameters in Table 1 (the rest is default).
2. Click on ‘Execute’ and wait until the processed is done.
3. Check out the output which is a collapsed sequenced in ARF format. For more infromation about this format, refer the documentation at the bottom of the miRDeep2 Mapper tool.

**Table 1:** miRDeep2 Mapper parameters.

| Options | Parameters |
| --- | --- |
| 'Deep sequencing reads' | choose your trimmed RNA-seq data |
| 'Remove reads with non-standard nucleotides' | 'Yes' |
| 'Select a reference genome' | choose your organism of reference |

#### 2.2.2 Getting the miRNA reference

Prior of mapping our collapsed read onto the known miRNAs, we first need to obtained our miRNAs from databases available online such as miRBase^8^ and miRTarBase^9^. Here we demonstrated how to obtain the miRNAs sequence from miRBase.

1. Go to the miRBase website (https://mirbase.org/) and click ‘Browse’ tab.
2. Choose your organism of interest e.g: Human. To see the full list of available organisms, click on ‘Expand All’.
3. A new page will be opened containing the summary of all available miRNAs of that organism. Go to the bottom of the page, select sequence type to ‘Stem-loop sequence’ and output format is ‘Unaligned FASTA format’.
4. Click ‘Fetch sequences’ and your reference sequence will be downloaded.
5. Repeat step 3 and 4 for ‘Mature sequence’.
6. Upload both of the sequence into your Galaxy history.

#### 2.2.3 Mapping to the miRNA using miRDeep2 Quantifier

miRDeep2 Quantifier will counts the amount of miRNAs available from our collapsed RNA-seq reads prepared prviously in 2.1 by mapping to the miRNAs references from step 2.2.

1. On the Galaxy’s tools panel, search miRDeep2 and click ‘miRDeep2 Quantifier’.
2. Set the following parameters according to the Table 2 (the rest is default, but also consider adjustment of other parameters according to your goal of experiment).
3. Click on ‘Execute’ and wait until the job is done
4. miRDeep2 Quantifier will output the list of mapped miRNAs including the counts and HTML report. Opened up these files to check out the read counts of miRNAs detected from your RNA-seq data.

**Table 2:** miRDeep2 Quantifier parameters.

| Options | Parameters |
| --- | --- |
| 'Collapsed deep sequencing reads' | the output from miRDeep2 Mapper |
| 'Precursor sequences' | the stem loop fasta file from miRBase |
| 'Mature miRNA sequences' | the mature fasta file from miRBase |
| 'Search in species' | choose to search only in specific species, otherwise it will search in all available species |
| 'Sort reads by sample in PDF' | 'Yes', will allow the highest read to be in the top |

#### 2.2.4 Compiling the miRNAs counts

If you have more than one replicates, or samples with different treatments, you might want to compile the whole read miRNAs counts from the miRDeep2 Quantifier results into one single table. Here we demonstrate step by step how to do that.

##### 2.2.4.1 Formatting the results

Figure 2 is the graphical resprentative of this step to format our obtained data into tab-seperated value files that can be uploaded into Galaxy.

**Figure 2:**
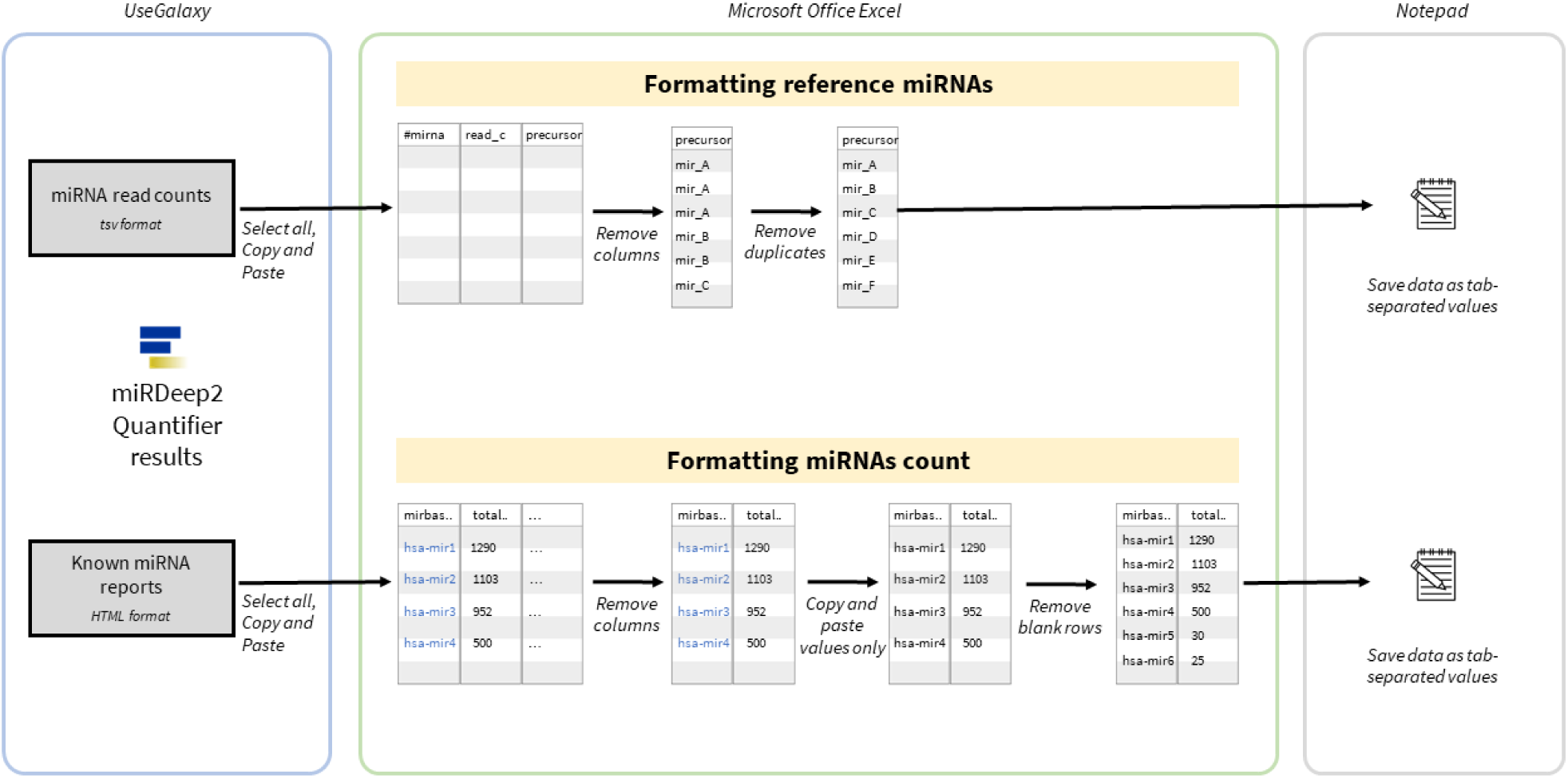
Overview of the data compiling step.

1. First we need to obtained the whole miRNAs reference that served as unique column in our table. Open one of the output table from miRDeep2 Quantifier results, right click and ‘Select all’, copy and paste into the spreadsheet (Microsoft Office Excel).
2. In the spreadsheet, remove other columns and keep the ‘precursor’ column.
3. To remove duplicates, at the ‘Data’ tab click on ‘Remove Duplicates’, select the column containing the miRNAs list and press ‘OK’. Make sure the first row contains the header named ‘precursors’ or you can rename if necessary. Save the file as ‘tab-separated values’ format and named it *miRNAs_ref*.*txt*.
4. Go to miRDeep2 Quantifier HTML report results in Galaxy, copy the whole table and paste into the new spreadsheet. Remove other columns and keep the first two columns which are ‘miRBase precursor ID’ and ‘total read count’. To make these as a header, remove other top rows.
5. Now we need to reformat into a clean table. Select the two columns and paste only the values into the next next column. Remove the original HTML formatted columns.
6. To remove the empty cells, select both columns and in the ‘Data’ tab click filter. Filter only the blanks on one of the column (check ‘Blanks’), select all the blank rows and delete; a pop out window will ask to delete the entire row, click ‘Yes’. Then set back to filter to show all the data.
7. To adjust the columns width (optional step), go to the ‘Home’ tab, click on ‘Format’ and ‘Autofit Row Heights’. Repeat the same for the ‘Autofit Column Width’.
8. Now we has a clean table with miRNAs precursors and read counts. Save the files as ‘tab-separated values’ format and named it *treatment_1*.*txt*. Consider also saving as an excel format for future editing.
9. Repeat the step 4 - 8 for other treatments. Alternatively you can set formatting of all treatments at the same time by compiling all data in one worksheet, and select the whole sheets at the same time before formatting. However saving need to be done one by one manually for each sheets.
10. Upload all of the saved files into the Galaxy, and set the format as ‘tabular’. Check each uploaded files to make sure there are no empty columns in between or at the end. If there are empty columns, remove it using ‘Cut columns from a table’ tool and use the edited file for downstream processing.

##### 2.2.4.2 Compiling the results in galaxy

We need to combine all the files into one single table count so that it would be easier for further analysis. Alternatively, spreadsheet can be use to combine all the tables and automatically fill the zero count miRNAs with 0. Here we demonstrate a galaxy tool to achieve the same result but with less computational burden to process the long spreadsheet.

1. Open up Galaxy and search ‘Join two files’ in the tool panel.
2. Select the miRNAs reference data (*miRNAs_ref*.*txt*) as the ‘1st file’ file to join and set ‘Column to use from 1st file’ to 1. Choose another data (*treatment_1*.*txt*) and set to ‘Column to use from 2nd file’ to 1. The other parameters are as is Table 3.
3. The output will contained the list of miRNAs from miRBase, and the read counts from your treatment. From this output, reapeat the step to combine other data to create one summary table of our miRNAs read counts. Now the data are ready for futher downstream processing.

**Table 3:** Join Two Files parameters.

| Options | Parameters |
| --- | --- |
| 'Output lines appearing in' | 'all lines [-a 1 -a 2]' |
| 'First line is a header line' | 'Yes' |
| 'Value to put in unpaired (empty) fields' | 0, this will fill in miRNAs with empty counts automatically |

## 3 Results & Discussion

### 3.1 Expected results

There are at least two main files created and processed by the user that are processed inside the local machine (miRNAs refernce and miRNAs count) and the most important output from this whole procedure is the combined read counts created using Galaxy’s Join Two Files tool. From step 2.2.4.1, the formatted clean miRNAs precursors should looks like Table 4 in tab-seperated value (.*txt* format). Note that different organism has different miRNA precursors. Meanwhile for the read count data, it should be similar with additional ‘count’ column according to the identified miRNAs.

**Table 4:** The top 10 results of miRNAs precursors processed from human miRNAs reference in the miRDeep2 Quantifier result.

| miR_precursors |
| --- |
| hsa-let-7a-1 |
| hsa-let-7a-2 |
| hsa-let-7a-3 |
| hsa-let-7b |
| hsa-let-7c |
| hsa-let-7d |
| hsa-let-7e |
| hsa-let-7f-1 |
| hsa-let-7f-2 |
| hsa-let-7g |

To demonstrate the real world data analysis, we analysed deposited exosomal NK-cell RNA-seq data available from GEO repository with GEO accession number GSE150342^10^. Following the procedure described above, the final output should looks like Table 5 with three of the experimental duplicates of IL15 treatment. When all these replicates data were combined using the Galaxy’s Join two file tool, the empty miRNAs counts will automatically set to 0 as set by our parameter.

**Table 5:** Results showing top 15 miRNAs count processed using our protocol from sample control group and IL15 treatment with 3 replicates each.

| miR_precursors | Control_1 | Control_2 | Control_3 | IL15_1 | IL15_2 | IL15_3 |
| --- | --- | --- | --- | --- | --- | --- |
| hsa-let-7a-1 | 29 | 12 | 54 | 87 | 108 | 233 |
| hsa-let-7a-2 | 29 | 11 | 54 | 86 | 109 | 233 |
| hsa-let-7a-3 | 29 | 12 | 54 | 86 | 107 | 231 |
| hsa-let-7b | 0 | 0 | 0 | 0 | 0 | 0 |
| hsa-let-7c | 4 | 0 | 0 | 11 | 0 | 0 |
| hsa-let-7d | 12 | 9 | 24 | 30 | 92 | 149 |
| hsa-let-7e | 9 | 0 | 6 | 20 | 2 | 4 |
| hsa-let-7f-1 | 16 | 7 | 33 | 27 | 20 | 57 |
| hsa-let-7f-2 | 15 | 5 | 28 | 24 | 20 | 61 |
| hsa-let-7g | 21 | 14 | 35 | 29 | 40 | 121 |
| hsa-let-7i | 19 | 10 | 41 | 39 | 53 | 182 |
| hsa-mir-100 | 17 | 0 | 1 | 37 | 0 | 0 |
| hsa-mir-101-1 | 41 | 4 | 15 | 58 | 6 | 1 |
| hsa-mir-101-2 | 41 | 2 | 15 | 57 | 6 | 1 |
| hsa-mir-10394 | 0 | 0 | 0 | 1 | 1 | 0 |

### 3.2 Further analysis

To demonstrate the further analysis of the data obtained using our method, RNA-seq count from Table 5 were uploaded into Galaxy and were performed differential analysis using limma tool^11^. Data input was set to single count matrix, performed using limma-voom differential expression method, and the other parameters were set accordingly to the data table (factor, groups and contrast). Limma tool allows comparisons (contrast) between many groups and their outputs can be downloaded into user’s machine. The result of this analysis were summarised in the Figure 3.

**Figure 3:**
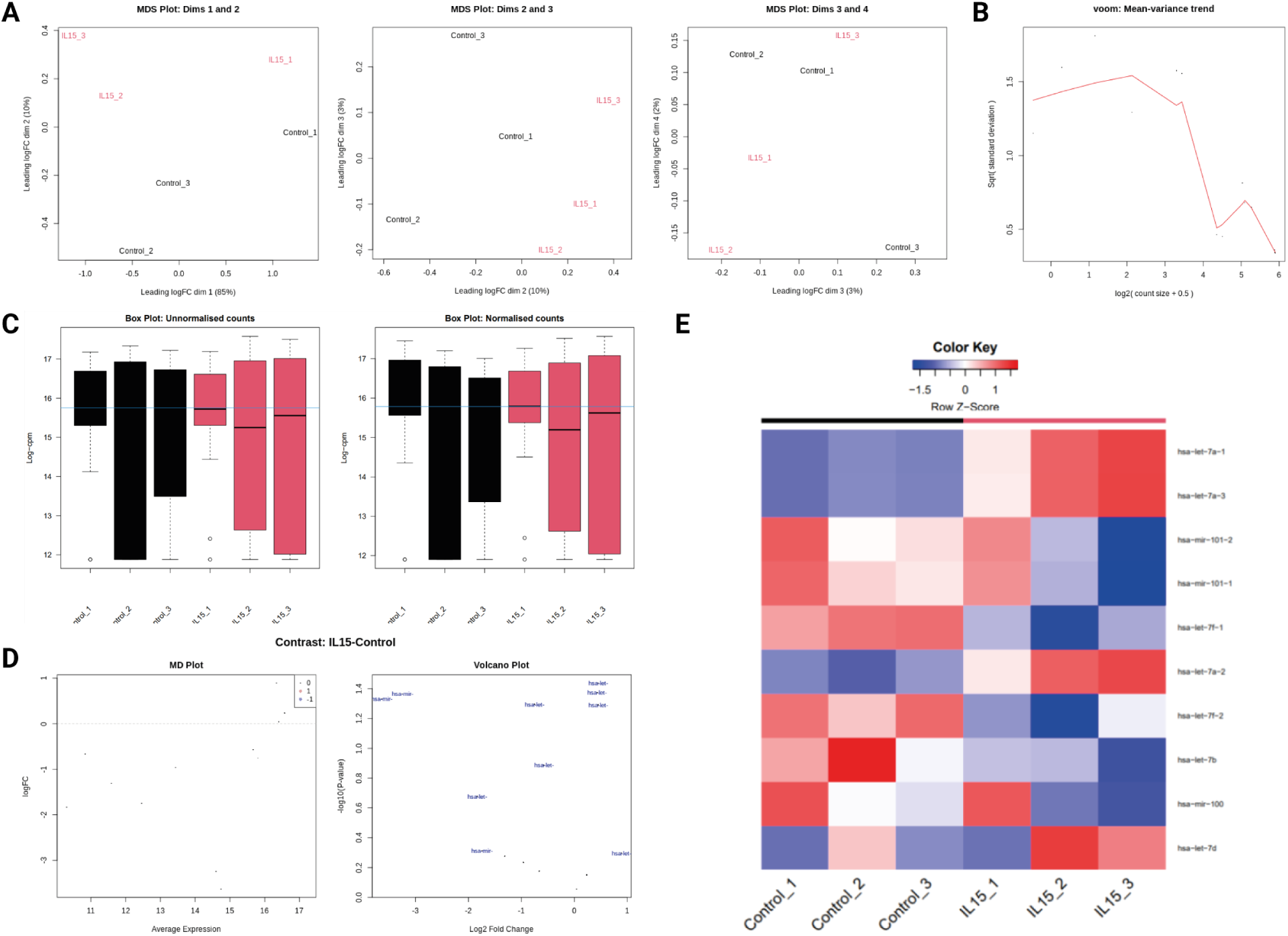
‘Differential analysis RNA-seq miRNAs count output from miRDeep2 of exosomal NK-cells with control and IL15 treatment using limma tool in Galaxy. A) The MDS plot at different dimensions: 85%, 10% and 3%. B) Mean variance trend of limma-voom. C) The boxplot of miRNAs counts before normalisation (unnomarlised) and after (normalised). D) The contrast set to compare IL15 and control group showing the MD plot and volcano plot of the two groups. E) The heatmap diagram of top 10 miRNAs showing the differential comparison of IL15 and control group in each samples.

## 4 Conclusion

Here we demonstrate our method to compiled and analysed the RNA-seq count data which is the output from miRDeep2 analysis. The proposed method, currently ultilised the scalability of Microsoft Office Excel and Join two file Galaxy’s tool that would allow user to process large input data which minimal user’s interference to minimise unwanted errors. We also tested the method for differential analysis using limma tool in Galaxy. Additionally, other differential analysis software such as edgeR and Degust could also use the same method with slight modification according to each tool’s input.

